# Denaturing mass photometry for straightforward optimization of protein-protein cross-linking reactions at single-molecule level

**DOI:** 10.1101/2023.05.30.542861

**Authors:** Hugo Gizardin-Fredon, Paulo E. Santo, Marie-Eve Chagot, Bruno Charpentier, Tiago M. Bandeiras, Xavier Manival, Oscar Hernandez-Alba, Sarah Cianférani

## Abstract

Mass photometry (MP) is a versatile, fast and low sample-consuming biophysical technique that gained interest in structural biology to study noncovalent assemblies in native conditions. We report here on a novel method to perform MP analysis in denaturing conditions (dMP) and its application for fast, accurate and straightforward optimization of chemical reactions in cross-linking mass spectrometry (XL-MS) workflows. dMP consists in a robust 2-step protocol that ensures 95% of irreversible denaturation within only 5 min. The proposed single-molecule method clearly overcomes the limitations and outperforms gold standard SDS-PAGE, as illustrated on several biological complexes. dMP provides an unprecedented and unmatched in-solution quantification of all coexisting XL species, including sub-complexes and non-specific XL aggregates, along with identification of significantly higher numbers of XL dipeptides in MS. We anticipate single-molecule dMP to be a high-impact game-changer for the XL-MS community with the potential to leverage the quality and reliability of XL-MS datasets.

## MAIN

Characterization of protein-protein interactions (PPI) are essential for the understanding of biological processes that drive life in different organisms. Although technological breakthroughs in well-stablished atomic-resolution biophysical techniques such as X-ray crystallography (XRC)^1^, nuclear magnetic resonance (NMR)^2^, and electron microscopy (EM)^3^ have boosted structural biology, several limitations (requirement of highly purified systems, extensive sample preparation protocols, production of high-quality crystals or cryo-EM grids, large amount of sample material, etc.) still remain. To circumvent those limitations and for NMR/XRC/cryo-EM-resistant complexes (highly flexible, disordered, membrane solubilized, etc.), structural mass spectrometry approaches nicely complement the classical structural biology toolbox^4,5^. Structural MS is a generic term that encompasses a series of MS-based strategies adapted for the characterization of non-covalent assemblies. Among them, protein cross-linking followed by mass spectrometry (XL-MS)^6^ is a covalent labelling proteomics-based technique that has drastically progressed this last decade from *in vitro*^7^, to proteome-wide^8^ or even *in-situ* and *in-cellulo* applications^9–11^. For XL-MS, the native (multi)-protein complex is usually incubated with a chemical reagent (cross-linker) in order to stabilize complexes (both stable and labile or transient), which allows to take a snapshot of non-covalent interactions, spatial proximities and provides insights on tertiary/quaternary structure and PPIs. The first step of chemical XL is of utmost importance: before going to MS analysis, optimal XL conditions have to be determined, ranging from the selection of the optimal XL reagent to screening of best adapted XL conditions ensuring proper XL while not generating non-specific XL aggregates^7^. According to latest published community-wide guidelines for XL-MS^12^, denaturing sodium dodecyl sulfate–polyacrylamide gel electrophoresis (SDS-PAGE) is recommended to monitor and optimize XL conditions, allowing visualization of high molecular weight (MW) bands on the upper part of the gel corresponding to cross-linked species, and concomitant vanishing of the bands corresponding to individual lower MW free protein partners. Even if SDS-PAGE analysis is easy, low cost and available in all laboratories, it also has some limitations, among which being: i) low mass-resolution, ii) time consuming (gel casting, sample denaturation, gel migration, staining) ; iii) not suited for high-mass complexes visualization as they do not enter the gel.

Mass photometry (MP), that has been recently introduced as a single-molecule biophysical technique^13^ operating in native conditions (nMP), can be positioned as alternative or to complement native mass spectrometry (nMS) for “nMS-resistant” protein complexes. MP is based on the principle of the interferometric scattering microscopy (iSCAT) and uses the contrast originated as a result of the destructive interaction between the scattered light and reflected light of biomolecules in solution upon irradiation with a visible laser light^14–16^. As the contrast intensity linearly scales with the mass, MP can serve to estimate masses of biomolecules after proper calibration with reference molecules. MP offers several advantages: i) a fast (minute scale), straightforward analysis of samples in solution, ii) multiplexing and automation capabilities, iii) no extensive sample preparation (e.g. buffer exchange)^17^, iv) low sample quantity requirements (pM-Nm concentration range, compared to μM range for nMS), v) a broad mass range (30 kDa to 5 MDa)^17,18^ and iv) single molecule detection which enables to relatively quantify the detected populations leading to the possibility to estimate affinity constants in the nM-μM concentration range^19,20^. Although nMS mass accuracy is unmatched compared to MP, nMP has already emerged as a valuable asset to characterize highly heterogeneous complexes^21^, membrane proteins solubilized with different types of membrane mimics^18,22,23^, ribosomes^24^, and viral capsids^25,26^.

Despite its routine use to study noncovalent machineries, MP is scarcely reported for the characterization and quantitation of covalent cross-linked assemblies. MP has been used after glutaraldehyde cross-linking of molecular machineries to verify the enrichment of a targeted oligomer before electron microscopy experiments^21^. As nMP gains popularity in the structural biology community, it looks appealing for straightforward XL reaction optimization. In this context, we report on the development of a groundbreaking and robust single-molecule protocol to perform MP analysis in denaturing conditions (dMP). We first used reference protein complexes of increasing sizes and complexities as proof-of-concept and dMP performance assessment. We then evaluated our dMP protocol for XL reaction monitoring and benchmarked it against the reference gold standard denaturing SDS-PAGE gel method. Due to its single molecule detection capabilities, we demonstrate here that dMP can be proposed as a straightforward, and more precise alternative to conventional SDS-PAGE analysis to monitor and optimize XL reactions, as illustrated on a real-case study for optimization of R2SP XL reaction.

## RESULTS

### Development of a denaturing mass photometry (dMP) protocol

To develop a fast, efficient and non-reversible denaturation protocol while maintaining good quality MP signal intensities, we step-by-step optimized several parameters and first focused on the choice of the denaturing agent. We selected as potential denaturing agents urea and guanidine HCl, two well-known chaotropic agents that disrupt H-bonds, increase solubility of hydrophobic regions, and finally lead to the disruption of tertiary and quaternary structures^27–29^, along with the H_2_0/ACN/FA mix (50/50/1) classically used in intact MS mass measurement in denaturing conditions. The H_2_0/ACN/FA mix was discarded, as incompatible with stabilization of MP droplets (which might be due to modification of the droplet surface tension related to presence of organic solvent and acid). We next assessed the impact of urea/guanidine HCl solutions on the quality of MP signal using “protein-free” droplets (**Fig. 1A**) by monitoring three output indicators (signal-*Si*, sharpness-*Sh* and brightness-*Br*) reflecting the quality of the MP images/frames. *Si* translates the level of activity in each frame that can be due to either protein binding, or contaminants/salts/surfactants presence. *Si* values should be as low as possible and always <0.05 % to avoid extensive peak broadening^17^. *Sh* refers to the level of detail visible in each frame, which influences the aptitude to find and maintain the good focusing position, and should be as high as possible. Finally, *Br* characterizes the amount of light available in images: as low *Br* will cause peak broadening by increasing the noise, *Br* value has to be maximized. With the aim to reach similar MP measurements qualities as those reached in PBS droplets (*Si* ∼0.03 %, *Sh* ∼5 %, *Br* ∼73 %), we used the “buffer-free” focusing mode to directly analyze protein-free droplets containing decreasing concentrations of urea or guanidine HCl (from 5.4 to 0.4 M). Of note, focusing the image on the glass surface was tedious for droplets concentrations higher than 1.75 M urea or guanidine HCl. Independently of the denaturing agent, *Sh* slightly increases progressively from 1-3% at 5.4 M of urea/guanidine HCl to ≥ 5 % at lower concentrations (**Table S1**). Similarly, *Br* increases at lower denaturing agent concentrations with an optimal value of ∼67 % obtained at 0.8 M urea/0.4 M guanidine HCl. *Si* values were < 0.05 % for all tested concentrations lower than 5.4M. Altogether, our “protein-free” blank MP acquisitions allowed establishing the optimal concentrations of denaturing agents (< 0.8M of urea or guanidine/HCl) in the PBS droplet in order not to compromise MP measurements quality. As XL reactions are typically conducted in the nM-μM concentration range, an approximatively 10x dilution in Phosphate Buffer Saline (PBS) is preconized prior to dMP measurements. That means that the initial concentration of the chaotropic agent during the denaturing reaction can be set at 5.4 M of urea and 6 M of guanidine without overcoming the 0.8 M concentration limit in the final droplet.

**Figure 1.**
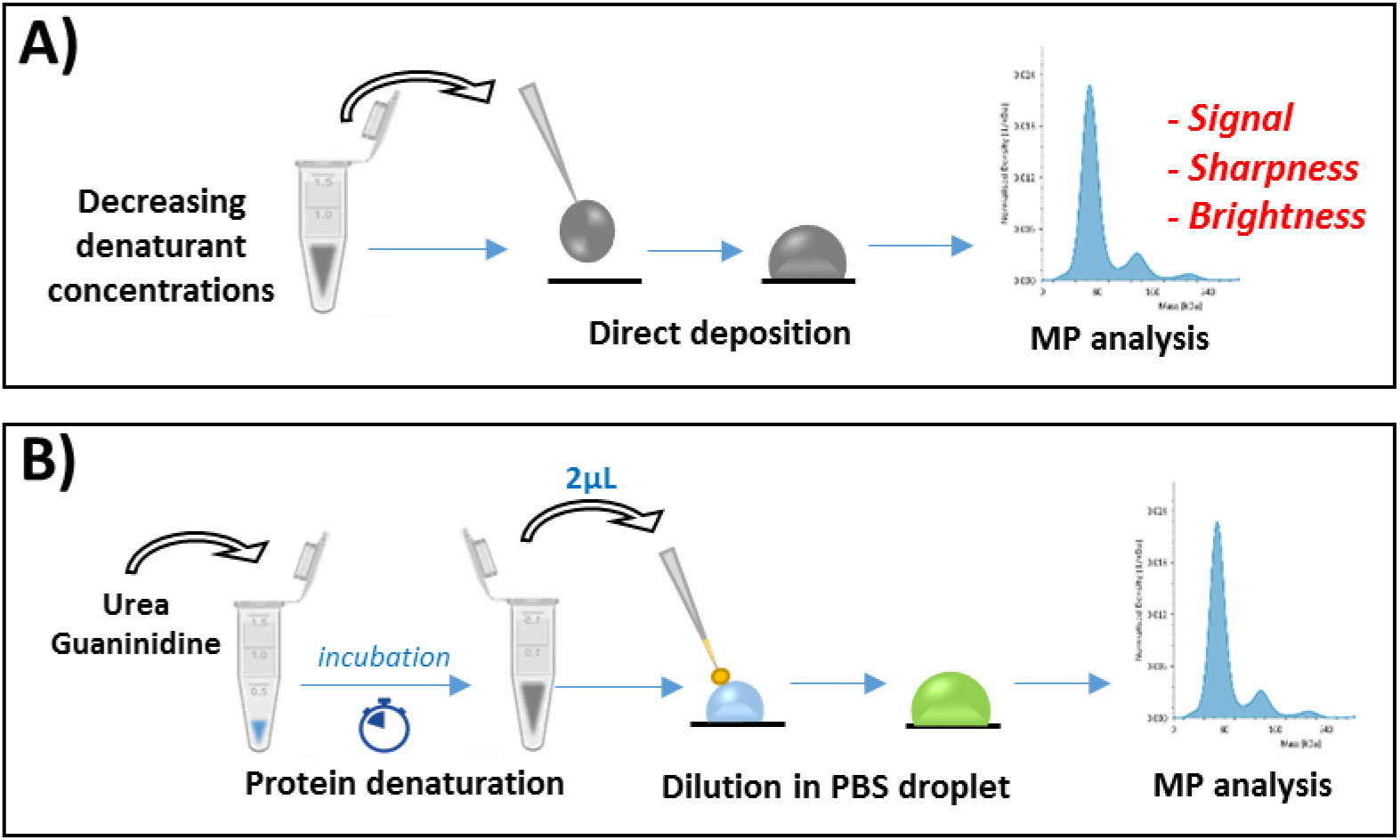
Schemes representing two MP-based assays used during dMP method development. **A)** Evaluation of of denaturing agents compatibility with MP measurements, **B)** Optimized general workflow for dMP analysis.

To further optimize our dMP protocol, we next used reference protein complexes (BSA, ADH, GLDH, 20S proteasome) either to assess mass precision, accuracy and peak broadening in dMP (BSA) or to further optimize the denaturation step (ADH, GLDH and 20S proteasome). After Gaussian-fitting of MP histograms, the mean mass (μ) and half-height peak width (2σ, FWHM) of the Gaussian fits were used to evaluate mass accuracy and peak broadening, respectively. Considering dMP triplicate measurements, the measured mass of BSA oligomers denatured in urea and guanidine HCl did not highlight any major differences compared to nMP measurements (**Table S2**). In addition, our denaturation protocol does not alter MP repeatability between replicates with SDs ≤ 3 kDa and ≤ 5 kDa for BSA monomers and dimers, respectively. Finally, denaturation only slightly affects peak broadening (FWHM between 8-10 kDa and 8-14 kDa, compared to 8 kDa in nMP for monomers and dimers, respectively), demonstrating that dMP measurements characteristics are similar to those obtained in classical nMP conditions.

In order to develop a fast denaturing protocol, we next optimized the duration of the denaturation step on proteins of increasing sizes and complexities (ADH, GLDH and 20S proteasome). Incubation in urea and guanidine HCl were carried out for 5 min to 16 h at room temperature (**Fig. 2**). After 5 min of urea denaturation, an almost complete denaturation is observed in dMP for all systems with > 95 % of the detected species being monomers (compared to 46 %, 19 % and 51 % of monomers for ADH, GLDH and 20S proteasome in nMP, respectively). Conversely, denaturation in guanidine HCl proved to be less efficient after 5 min for ADH (∼34 % monomers) while being equivalent for GLDH and 20S proteasome, suggesting urea as a more ubiquitous denaturing agent. In order to ensure that no protein refolding occurs in the PBS droplet prepared for dMP measurements, and in the timeframe of the analysis, we mimicked our optimized denaturing protocol in a tube (**Fig. S1 A)**). As expected, only GLDH monomer (61 ± 6 kDa) is detected even after 10 min in PBS, with perfectly superimposable dMP spectra (**Fig. S1 B)**), demonstrating that no significant protein refolding will occur within the timeframe of dMP analysis (typically 1 min).

**Figure 2.**
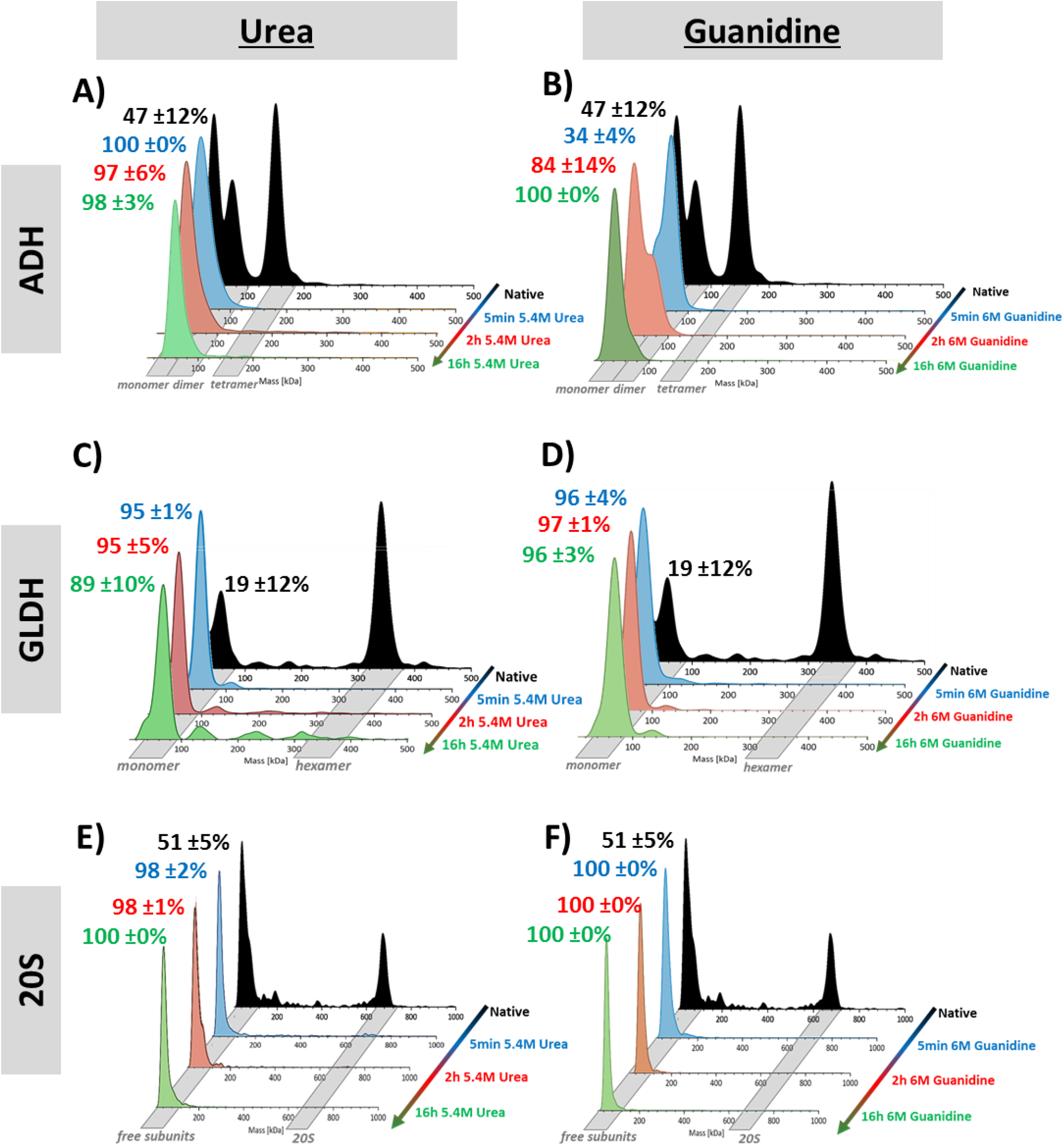
Optimization of denaturation step for dMP. Mass distributions, represented as probability density (KD), show the evolution of oligomeric states abundances. ADH denaturation in urea A) or in guanidine HCl **B)** ; GLDH denaturation in urea **C)** or in guanidine HCl **D)** ; 60S proteasome denaturation in urea **E)** or in guanidine HCl **F)**. For each protein and condition, the % of monomeric species are indicated with similar color-code as their respective time point.

To conclude, we have developed and optimized a fast (5 min), efficient (> 95 % denaturation) and non-reversible (in the timeframe of dMP measurements) denaturation protocol compatible with MP analysis, which will be further referred to as “denaturing MP protocol” (dMP). This workflow consists of a first step of denaturation (5 min in 5.4M urea) followed by dilution of 2 μL of the denatured sample to a 18 μL PBS droplet right before dMP analysis (**Fig. 1B**). Obtained dMP measurements quality along with mass accuracies and peak width were comparable with those obtained in classical nMP analysis.

### dMP outperforms SDS-PAGE gel analysis for XL reaction monitoring

We next benchmarked our dMP protocol against SDS-PAGE, the gold standard for XL reaction optimization, on our reference systems (ADH, GLDH, 20S proteasome), using the MS-cleavable cross-linker disuccinimidyl dibutyric urea (DSBU) as proof of concept. For a 25:1 DSBU:ADH ratio (**Fig. 3A**), monomers, dimers, trimers and tetramers are observed in dMP (50, 25, 14, 10 % of total counts, respectively), in good agreement with SDS-PAGE (**Fig. S2**). Tetrameric species are the most abundant from DSBU molar excesses ≥ 100 with up to 53 % of total counts. No non-specific high-mass aggregates were detected neither using SDS-PAGE nor dMP, both methods suggesting optimal conditions around 100:1 DSBU:ADH. For GLDH (**Fig. 3B**), dMP allows to visualize that increasing XL concentration progressively stabilize higher oligomeric states, with a reduction of intermediary sub-complexes (23 to 9 % from 100 and 400 DSBU molar excess) at the expense of the hexamer (57 to 79 % from 100 and 400 DSBU molar excess). At both 100:1 and 400:1 DSBU:GLDH ratios, only a small proportion of dodecamers (6 %) is formed, suggesting non-specific XL aggregates. Conversely, regardless the XL condition, only a broad band in the loading-well is observed on the SDS-PAGE gel at high masses, highlighting that GLDH high mass oligomers do not enter the gel, along with remaining free monomers still observed for all DSBU molar excesses. This impairs a proper assessment of stabilized oligomerization states using this technique. For cross-linked 20S proteasome (28 subunits), SDS-PAGE is even less useful, as only a slight decrease in the free subunits intensity can be seen upon cross-linker concentration increase (**Fig. 3C**). Additional masses between 100-200 kDa are also stabilized, but, in all cases, SDS-PAGE does not allow to visualize the ∼700 kDa 20S proteasome assembly. Conversely, dMP mass range revealed 20S proteasome covalent stabilization even at 25:1 DSBU:20S ratio (16 %) with a maximal 20S cross-linking at 400:1 DSBU:20S (25 %), while avoiding non-specific aggregation.

**Figure 3:**
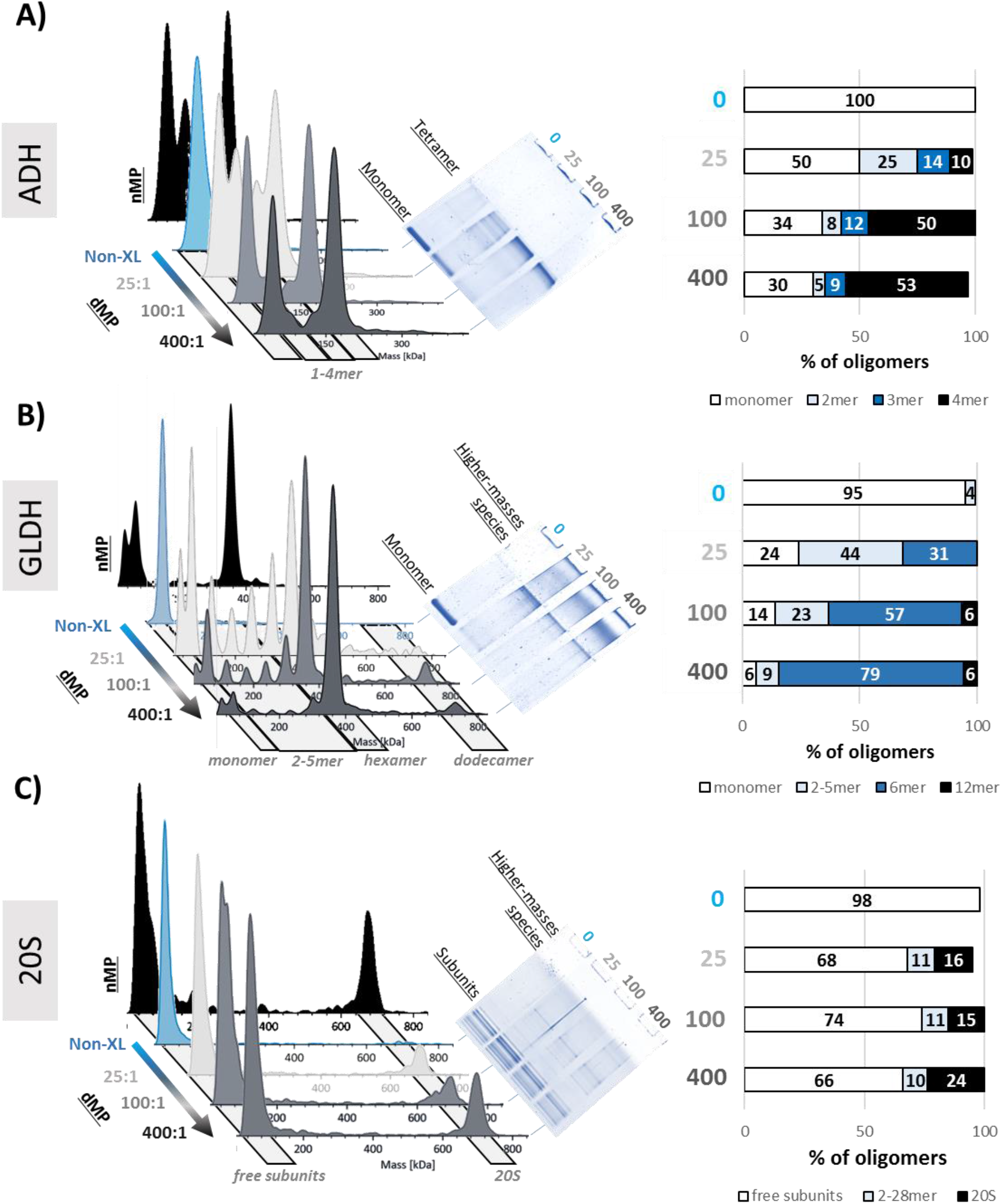
Benchmarking of dMP vs SDS-Page for cross-linking optimization. Mass distributions are represented as probability density (KD). Following samples were cross-linked with increasing molar excesses of DSBU: **A)** nMP of non-cross-linked ADH, dMP and SDS-PAGE results of ADH cross-linking. **B)** nMS of non-cross-linked GLDH, dMP and SDS-PAGE results of GLDH crosslinking **C)** nMS of non-cross-linked 20S proteasome, dMP and SDS-PAGE results of 20S proteasome crosslinking. Bar plots show calculated dMP-calculated percentages of oligomeric populations for each concentration of DSBU.

Thanks to its single molecule detection capabilities along with higher mass accuracies and increased mass range (30 kDa-5 MDa) compared to SDS-PAGE (10 kDa-300 kDa), dMP allowed fast in-solution mass measurements and relative quantification of all co-existing single molecules detected. This is of utmost advantage for high mass and high-heterogeneity samples for which SDS-PAGE clearly fails to give the needed output for a rational XL optimization. In addition, dMP highlights that the use of high XL reagent concentrations usually not recommended (400 molar excesses) only leads to a very limited amount of non-specific XL aggregates. Altogether dMP, that allows single-molecule sensitivity, proved to be more precise, quantitative and provides less arbitrary optimization than SDS-PAGE.

### dMP offers unmatched rapidity for straightforward quantitative screening of optimal XL conditions

We next evaluated the versatility of dMP to monitor XL reaction using a variety of chemical reagent : two MS-cleavable compounds, DSBU (disuccinimidyl dibutyric urea, linker size ∼ 12.5 Å) and DSAU (disuccinimidyl diacetic urea, linker size ∼7.7 Å) and the less-flexible IMAC-enrichable cross-linker PhoX (linker size ∼5.5 Å) that gains popularity for both *in vitro* and *in vivo* XL-MS studies^6,30^. For GLDH, obtained dMP mass distributions are similar for PhoX and DSAU cross-linking, but clearly differ from DSBU (**Fig. 4a**): while GLDH hexamers and monomers are detected as main components at 25:1 DSBU:GLDH ratio, monomers, dimers and trimers are formed in similar conditions with DSAU and PhoX. This trend is even more obvious at 100:1 and 400:1 cross-linker:GLDH ratios, with an almost complete stabilization of GLDH hexamers using DSBU when only low abundant hexameric species are detected with PhoX and DSAU. Similar behaviors were also observed in dMP profiles of cross-linked ADH and 20S proteasome, with a significantly higher stabilization of intact complexes with DSBU (**Fig. S3 A)** and **B)**). To go deeper into cross-linker comparison, we defined two metrics for rational and quick comparison of different XL conditions (reagent, reaction time, pH etc): i) the global inter-protein XL reaction efficiency Eff_XL_ (Material and method **Eq. 1**), as an indicator of the relative amount of inter-protein XL-stabilized species (all cross-linked stabilized species except monomers); and ii) a XL-stabilization factor SF_XL_ (Materials and Methods **Eq. 2**), as an estimate of the amount of “expected XL-stabilized complex”.

**Figure 4:**
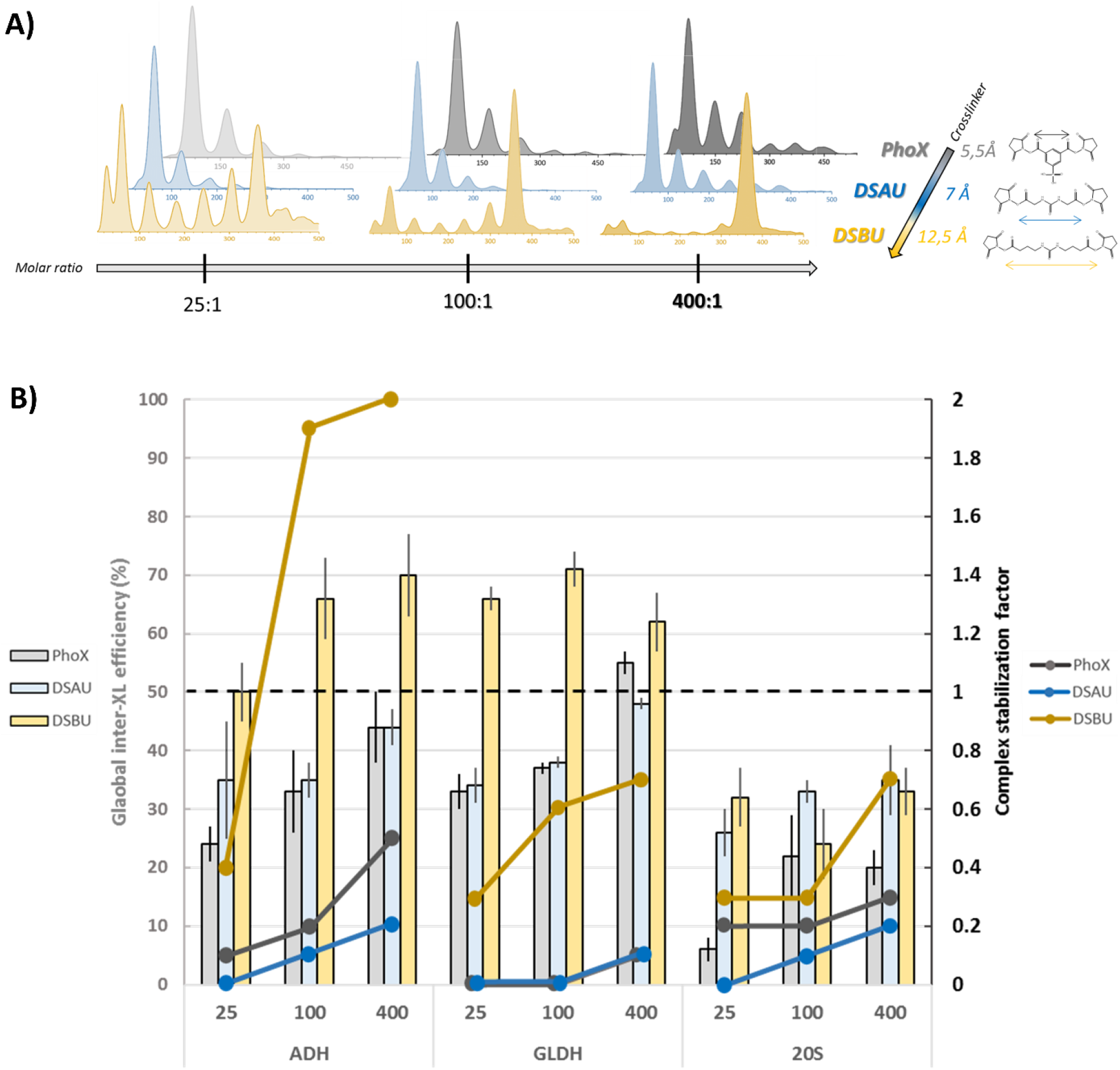
dMP screening of cross-linking conditions on ADH, GLDH and 20 proteasome. **A)** *Effect of cross-linking reagent (size, flexibility) on oligomeric states stabilized, measured in dMP*: presented results are probability densities (KD) of GLDH samples cross-linked with increasing molar ratios of PhoX, DSAU, DSBU. **B)** *dMP-based quantitative results of XL condition screening for ADH, GLDH and 20S complexes*. Bar charts represent the dMP-calculated global inter-XL efficiency (% ± SD) for each complex and XL condition (25/100/400:1 cross-linker:complex molar ratio). Plain dots represent the complex stabilization factor for each complex and XL condition. The black dash line corresponds to the stabilization factor value of 1 indicating a complex abundance similar to the native sample.

Eff_XL_ estimation is thus an interesting indicator for XL conditions screening but lacks in precision as it takes into account both expected specific and non-specific XL species, which is not the case of SF_XL_ that focuses on one species. The SF_XL_ factor should be maximized to get close to a value of 1 for a complete XL-stabilization of the non-covalent complex : SF_XL_ values > 1 means stabilization of expected complex’s abundance along with a decrease in generation of transient XL-sub-complexes assemblies, that might occur over the XL reaction time (30 min to 1 hour). For homo-oligomeric samples (ADH and GLDH), DSBU showed much higher Eff_XL_ values (60-70 %) compared to DSAU (∼45 %-48 %) and PhoX (45-55 %) (**Fig. 4B**, bar charts), suggesting overall better XL efficiencies for DSBU. For the hetero-multiprotein 20S proteasome, trends are different: i) the Eff_XL_ is decreased (max. ∼35 %) and ii) DSAU and DSBU are the most potent cross-linkers over PhoX. For all reference systems, SF_XL_ values : i) increase as a function of cross-linker:complex ratio, suggesting increased stabilization of expected covalent assemblies at 400:1 cross-linker:complex ratio and ii) are higher for DSBU compared to PhoX and DSAU that exhibit the lowest SF_XL_ values (**Fig. 4B**, solid dots). Low SF_XL_ values combined with good Eff_XL_ (26-48 %) translate DSAU abilities to generate more sub-complexes but low amounts of expected XL-stabilized ADH/GLDH/20S proteasome tetramers/hexamers/28-mer. Of note, it appears that SF_XL_ values obtained for ADH and GLDH tend to plateau with increasing DSBU molar excess, which is not the case for the 20S proteasome, constituted of a higher number of subunits (28) compared to AHD (4) and GLDH (6).

Finally, we applied dMP for *ab initio* fine-tuning of R2ΔIIS’P’ (here called R2SP) cross-linking, ∼540kDa (**Fig. S4**) complex constituted of RuvB-Like1 without the DII_ext_ (R1ΔDII, R1) and RuvB-Like2 without the DII_ext_ (R2ΔDII, R2)^31^, SPAG1_622-926_ (S’) and PIH1D2_231-315_ (P’)^32,33^. We compared 4 amine-reactive cross-linkers^6^, including PhoX (5.5 Å), DSAU (7Å), DSSO (10.3 Å), DSBU (12.5 Å), each at 5 different molar excesses (25/50/100/200/400) plus the control non-XL sample (**see Fig. S5 for SDS-PAGE**). This 24-conditions screen was realized with unprecedented rapidity (∼1.5 hours), including calibration and dMP triplicates measurements. Despite a dMP a progressive stabilization of sub-complexes with increasing cross-linker concentration of PhoX (Eff_XL_ ∼31-35%, max. SF_XL_ ∼0.1) and DSAU (Eff_XL_ ∼31-35 %, SF_XL_∼0, **Fig. 5E**), none of these reagents are able to significantly stabilize the ∼540 kDa R2SP detected in nMP (**Fig. 5A and 5B**). In contrary, both DSSO (Eff_XL_ ∼31-35 %, max. SF_XL_ ∼0.4) and DSBU (Eff_XL_ ∼31-35 %, max. SF_XL_ ∼0.7) at high molar excesses (> 100:1 DSSO/DSBU:R2SP) yield to stabilization of covalent ∼540 kDa R2SP, without significant non-specific aggregate formation or sub-complex stabilization (**Fig. 5C-E**). From dMP, a 400:1 DSBU:R2SP ratio (max. Eff_XL_ ∼77 ± 6 %) would be selected as optimal XL condition for further XL-MS analysis (**Table S3**), which confirmed identification and validation of a higher proportions of inter-XL peptides (from 45 % for 25:1 DSBU:R2SP to 55 % for 400:1 DSBU:R2SP).

**Figure 5:**
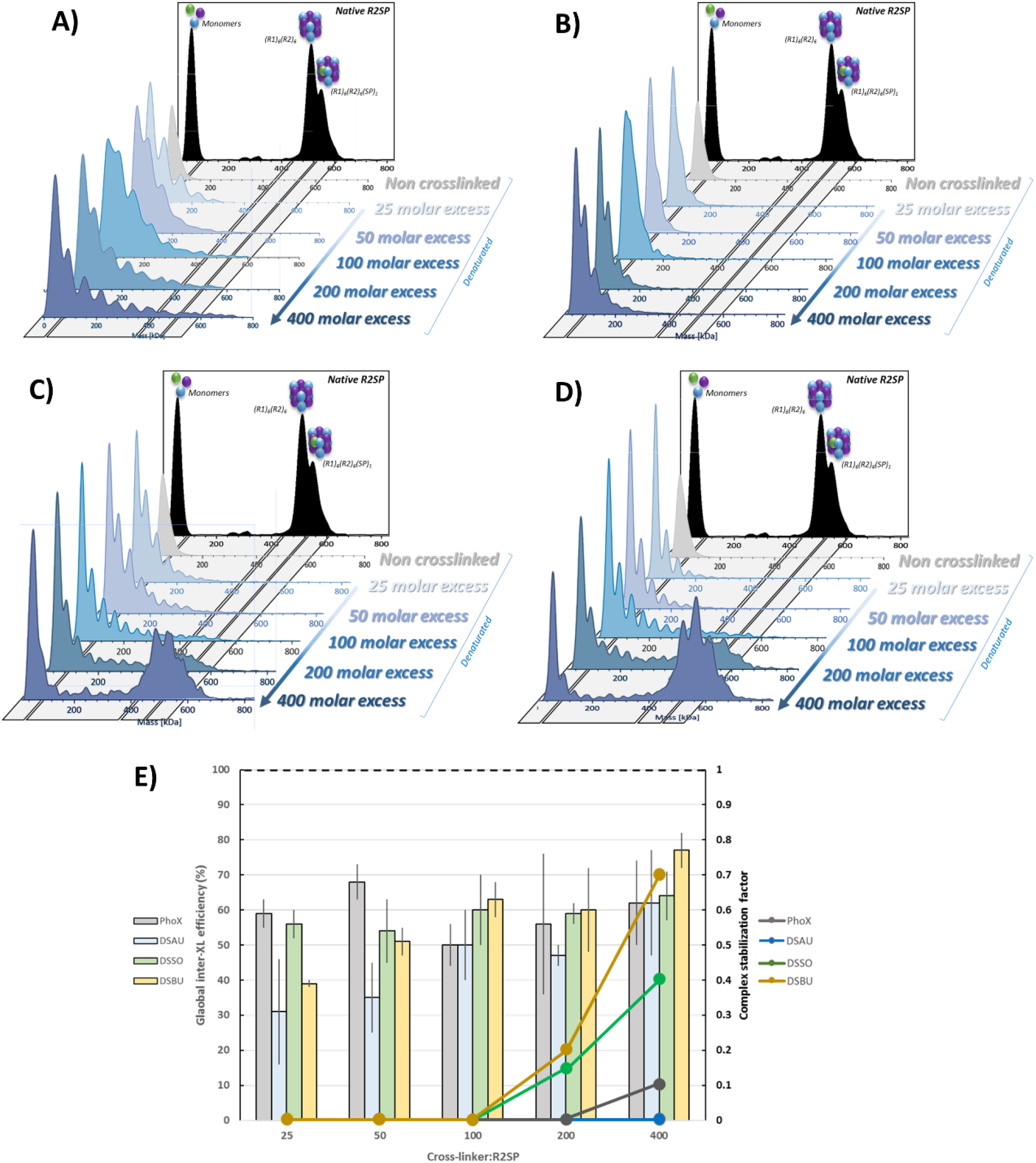
dMP results of XL reaction optimization for R2SP complex. dMP profiles of R2SP cross-linked with increasing molar excesses of **A)** PhoX, **B)** DSAU, **C)** DSSO, **D)** DSBU. **E)** Quantitative results of R2SP XL optimization. Bar charts represent the dMP-calculated global inter-XL efficiency (% ± SD) for each complex and XL condition. Plain dots represent the complex stabilization factor for each complex and XL condition. The black dash line corresponds to the stabilization factor value of *1* indicating a complex abundance similar to the native sample.

Altogether, our results demonstrate that dMP affords unmatched performances for fast and rational screening for optimal XL conditions (e.g. reagent, reaction length, temperature, buffer, etc.). Our dMP results highlight that DSBU appears to outperform DSAU, PhoX, and even DSSO for inter-protein XL on our biological systems. This reagent provides the best compromise between stabilization of the covalent-XL-assembly of interest (SF_XL_ values close to 1 while minimizing covalent sub-complexes stabilization or non-specific aggregate formation.

## DISCUSSION

We report here on the development of a MP protocol allowing analysis in denaturing conditions, thereby addressing the demand for versatile tools for XL reaction monitoring to better understand biologically relevant PPIs. The dMP approach consists in a fast and efficient denaturing protocol that does not alter the quality of the MP measurement, resulting in > 95 % denaturation of the sample within 5 minutes (using urea 5.4 M followed by dilution of 2 μL in a classical PBS droplet). The applicability of the dMP protocol is illustrated to fine tune XL reaction conditions. Our results highlight that dMP outperforms SDS-PAGE analysis in XL-MS workflows, allowing faster (few minutes) and more precise mass measurements along with relative quantification of all the different single-molecule resolved cross-linked species. We propose dMP as an empowered technique alternative to SDS-PAGE analysis to screening for best XL conditions before MS analysis. Several lessons can be learned from this pilot dMP study, as our results challenge a number of XL-MS well-established rules: i) chemical XL reactions are not necessarily low efficiency; ii) a large molar excess of cross-linkers does not necessarily generate artefactual XL non-specific aggregates; iii) size and flexibility of the chemical reagents drive the stabilization of XL-species and iv) MS-cleavable DSBU and DSSO might be best adapted compared to PhoX or DSAU for *in vitro* intra-protein XL workflows.

Thanks to its unique single-molecule detection capabilities, dMP allowed addressing the question of the “low” yield of the chemical XL reaction (< 10 %)^30^, often correlated to the low number of identified XL dipeptides (low abundant XL dipeptides compared to linear non-XL peptides). Our dMP results reveal that significant stabilization efficiency of covalent XL-assemblies (> 50 %) can be achieved and easily approximated through SF_XL_ calculations (target SF_XL_ ∼1). Another fundamental point raised by our pilot dMP study concerns the dogma in the XL-MS community to perform XL reactions at a relatively low excess of cross-linkers (50-200x), to avoid the generation of non-specific XL resulting from “forced pairing” of distal amino acid sequences that might not be biologically relevant. Surprisingly, our dMP results rather highlight that: i) low molar excess of XL reagents (100-200x) would rather stabilize sub-complexes while ii) a high excess (400x) would be required to stabilize expected covalent XL assemblies as a whole.

dMP results also strengthen the importance of the XL reagent choice for successful XL-MS analysis. The choice of the XL reagent is guided by several properties ranging from the amino acids to be targeted (homo-versus hetero-reagents), the length of the XL reagent (that defines the reachable distances) but also by the chance to identify XL peptides by MS/MS (MS cleavable versus non cleavable reagents). MS-cleavable cross-linkers (such as DSSO, DSAU and DSBU) along with IMAC-enrichable reagents (PhoX) are mostly used because they allow more straightforward MS/MS identification of the XL-peptides. Our dMP results suggest that inter-protein XL efficiency/stabilization depends on the length and flexibility of the spacer arm. DSBU thus provides better XL efficiencies/stabilizations (Eff_XL_, SF_XL_) than other tested reagents, which is consistent with the 12.5 Å highly flexible DSBU increased potential to access a higher number of potential K-K pairs over a wider distance range compared to less flexible and smaller sized DSAU and PhoX, in agreement with published data^34–36^. Conversely, less flexible XL reagents with smaller sized spacer arms might rather lead to stabilization of sub-complexes instead of “expected native complexes”. PhoX could finally be more suited for intra-protein XL.

To conclude, the developed single molecule dMP strategy not only provides an unmatched increase in the speed and quality of screening for optimal XL conditions, but also provides identification and relative quantification of all single molecule coexisting cross-linked species, including sub-complexes and non-specific XL aggregates, previously not clearly identified by SDS-PAGE. dMP uniquely provides evidence for the rational quantitative selection of best-adapted chemical XL conditions based on XL stabilization/efficiency indicators. Altogether, we anticipate single-molecule dMP to be a high-impact game-changer go-to method to leverage the quality and reliability of XL-MS datasets by providing direct snapshots of all coexisting species.

## MATERIALS AND METHODS

### Stocks, reagents, and instruments

The complete list of reagents and instruments used in this study are listed in Materials section in the Supplementary Information.

### Sample preparation and cross-linking reactions

Bovine Serum Albumin (BSA, Sigma, Saint-Louis, USA), Alcohol dehydrogenase from baker’s yeast (ADH, Sigma, Saint-Louis, USA) and L-Glutamate dehydrogenase from bovine liver (GLDH, Sigma, Saint-Louis, USA) were diluted to 1mg/mL in Gibco^TM^ phosphate buffer saline (PBS, Life technologies Corporation, NY, USA), pH 7.4. Human 20S proteasome (20S, South Bay Bio, San Jose, USA) was diluted to 1mg/mL in 50 mM HEPES, 100 mM NaCl, pH 7.4 prior to cross-linking.

R2SP complex complex was formed by incubating pure RuvBL1/RuvBL2 with excess pure SPAG1/PIH1D2 at a ratio of 1:4, respectively. The mixed complexes were incubated over night at 4 °C and the formed R2SP complex was separated from excess SPAG1/PIH1D2 using a Superose 6 16/60 XK (GE Healthcare), see Supplementary Information for detailed protocol. It was diluted to 1mg/mL in 20 mM Hepes, 150 mM NaCl, pH 8^31^ prior to cross-linking. For cross-linking (XL) reactions, aliquots of XL reagents were freshly diluted in DMSO. Following reagents were used: PhoX (Disuccinimidyl Phenyl Phosphonic Acid, Bruker); DSAU (Disuccinimidyl diacetic urea, CF Plus Chemicals, Brno-Řečkovice, Czech Republic); DSSO (Disuccinimidyl sulfoxide, Thermo Fischer Scientific, Rockford, IL, USA); DSBU (Disuccinimidyl dibutyric urea, CF Plus Chemicals, Brno-Řečkovice, Czech Republic). BSA, ADH, GLDH and 20S samples were each split in six aliquots and incubated with 25, 100, or 400 molar excess of each reagent. R2SP stock solution was split into aliquots subsequently reacted with PhoX, DSAU, DSSO, DSBU at molar excesses of 25, 50, 100, 200, 400.

XL reactions were carried for all samples at room temperature (18°C) for 45 min, and quenched with Tris HCl (15 mM final concentration) for 20 min. A 1.5 μg aliquot of each non-XL control and XL sample was kept for SDS-PAGE migration.

### SDS-PAGE separation of cross-linked samples

All cross-linked proteins and complexes were migrated on in-house 12% acrylamide denaturing SDS-PAGE gels (1.5 mm thickness). Volume corresponding to 1.5 μg of each XL sample (and non-XL controls) was diluted (1:1) with 2x concentrated Læmmli buffer (4% SDS, 20% glycerol, 10% 2-mercaptoethanol, 0.01% bromphenol blue and 0.125 M Tris HCl) and incubated 5 min at 95°C. After sample loading, gels were migrated at 50 V for 20 min, 100 V until the 2/3 of the gel and 120 V until the end. After migration, gels were fixated for 20 min (3 % phosphoric acid, 50 % ethanol), washed 3×20min with milli-Q water and stained overnight with Coomasie Brillant Blue (G250, Sigma, Saint-Louis, USA). They were finally rinced 3×20 min with milli-Q water.

#### Mass photometry measurements

MP measurements were performed with a TWO^MP^ (Refeyn Ltd, Oxford, UK) at room temperature (18 °C). Microscope slides (24×50 mm, 170±5 μm, No. 1.5H, Paul Marienfeld GmbH & Co. KG, Germany) were cleaned with milli-Q water, isopropanol, milli-Q water and dried with a clean nitrogen stream. Six-well reusable silicone gaskets (CultureWell^TM^, 50-3 mm DIA × 1mm Depth, 3-10 μL, Grace Bio-Labs, Inc., Oregon, USA) were carefully cut and assembled on the cover slide center. After being placed in the mass photometer and before each acquisition, a 18 μL droplet of PBS was put in a well to enable focusing on the glass surface.

#### Contrast-to-mass calibration

To allow MP mass measurements, contrast-to-mass calibration was performed twice a day by measuring a mix of Bovine Serum Albumin (66 kDa), Bevacizumab (149 kDa), and Glutamate Dehydrogenase (318 kDa) in PBS buffer, pH 7.4. The distributions of scattering events (given as contrast) were Gaussian-fitted using DiscoverMP (Fig. S6 A)). Contrasts values are converted into masses using linear relation between the contrast and the mass of the binding object. Calibrations were accepted for R2>0.995 (Fig. S6 B).

#### Native MP (nMP)

Samples were first diluted with their native buffer to 100-400 nM. Finally, 2 μL of the stock solution are finally drop-diluted and carefully mixed to 10-40 nM in a 18 μL PBS droplet^17^. Three movies of 3000 frames were recorded (60 s) for each sample using the AcquireMP software (Refeyn Ltd, Oxford, UK).

#### Denaturing MP (dMP)

Denaturing MP experiments were carried out by incubating first the samples to a protein concentration of 100-400 nM in 5.4 M Urea (Sigma, Saint-Louis, USA) or 6 M Guanidine (Sigma, Saint-Louis, USA). For non-crosslinked samples incubation times evaluated ranged from 5 min to 16 hours at room temperature (18°C). After incubation and right before MP measurements, 2 μL of the solution were quickly drop-diluted^18^ in an 18 μgL PBS droplet to 10-40 nM. All measurements were done immediately following the droplet dilution. For the final optimized dMP protocol, denaturation was done in Urea 5.4 M for 5 min.

#### MP Data processing

Data were processed using the DiscoverMP software (Refeyn Ltd, Oxford, UK). Obtained distribution histograms represent the number of counts per contrast value (or per mass after calibration). To obtain the average masses, peak width and number of counts for each mass distribution, a Gaussian fitting was performed by integrating each distributions at its half-height. Relative amounts of each oligomer were calculated using the number of counts under the Gaussian fit curve of each distribution. For figures, Kernel Density Estimate (KDE) was applied to transform the histogram into a curve.

#### Calculation of the global inter-protein cross-linking reaction efficiency (Eff_XL_)

Global inter-XL efficiency was calculated using number of counts after Gaussian fitting of each oligomeric state distribution (example of calculation in **Table S4**). This value represent the efficiency of XL reaction to stabilize inter-protein interactions, i.e. all oligomeric states > 1 remaining after denaturation. Inter-XL efficiency does not discriminate specific interactions of unspecific aggregation.

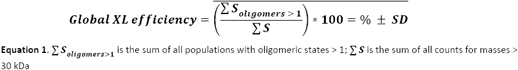

#### Calculation of the complex stabilization factor (SF_XL_)

The non-XL native measurements were used as a reference to obtain the proportion represented by the complex to be XL-stabilized in the sample. Then, we similarly calculated the proportion of this complex among total counts of the cross-linked denatured sample. Using these two values, the complex stabilization factor can be calculated (example of calculation in Table S5). This value expresses the amount of native complex that could effectively be XL-stabilized in XL samples (Eq. 2). A factor value of 1 correspond to the stabilization of all the native complex after XL reaction. Value > 1 expresses an enrichment of the complex upon XL reaction. Stabilization factor should be ideally ≥ 1.

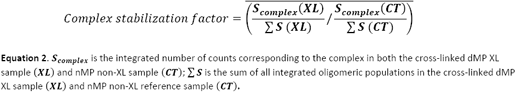

## Supporting information

supp data all

## DATA AVAILABILITY

XL-MS datasets are deposited in the ProteomeXchange Consortium via the PRIDE^37^ partner repository under the number PDX….

## ACKNOWLEDGMENTS

This work was supported by the CNRS, the Univserity of Strasbourg, the “Agence National de la Recherche” and the French Proteomics Infrastructure (ProFI; ANR-10-INBS-08-03). H.G.F. acknowledges the French Ministry for Education and Research for funding of his PhD. This work was funded by Fundação para a Ciência e Tecnologia/Ministério da Ciência, Tecnologia e Ensino Superior (FCT/MCTES, Portugal) through national funds to iNOVA4Health (UIDB/04462/2020 and UIDP/04462/2020) and the Associate Laboratory LS4FUTURE (LA/P/0087/2020).

## AUTHOR CONTRIBUTIONS

H.G.F performed the experiments. H.G.F., O.H, S.C conceived the study. S.C supervised the work. H.G.F., O.H, S.C wrote the manuscript. M.E.C. carried out the cloning of the domains of SPAG1 and PIH1D2. P.E.S. carried out the expression and the purification of R2SP complex. P.E.S, M.E.C., B.C., T.M.B. and X.M. proofread the manuscript.

## REFERENCES

1. Shi, Y. A Glimpse of Structural Biology through X-Ray Crystallography. Cell 159, 995–1014 (2014).

2. Markwick, P. R. L., Malliavin, T. & Nilges, M. Structural Biology by NMR: Structure, Dynamics, and Interactions. PLoS Comput. Biol. 4, e1000168 (2008).

3. Bai, X., McMullan, G. & Scheres, S. H. W. How cryo-EM is revolutionizing structural biology. Trends Biochem. Sci. 40, 49–57 (2015).

4. Britt, H. M., Cragnolini, T. & Thalassinos, K. Integration of Mass Spectrometry Data for Structural Biology. Chem. Rev. 122, 7952–7986 (2022).

5. Jooß, K., McGee, J. P. & Kelleher, N. L. Native Mass Spectrometry at the Convergence of Structural Biology and Compositional Proteomics. Acc. Chem. Res. 55, 1928–1937 (2022).

6. Piersimoni, L., Kastritis, P. L., Arlt, C. & Sinz, A. Cross-Linking Mass Spectrometry for Investigating Protein Conformations and Protein–Protein Interactions─A Method for All Seasons. Chem. Rev. acs.chemrev.1c00786 (2021) doi:10.1021/acs.chemrev.1c00786.

7. Chavez, J. D. et al.. Systems structural biology measurements by in vivo cross-linking with mass spectrometry. Nat. Protoc. 14, 2318–2343 (2019).

8. Klykov, O. et al.. Efficient and robust proteome-wide approaches for cross-linking mass spectrometry. Nat. Protoc. 13, 2964–2990 (2018).

9. Sinz, A. Crosslinking Mass Spectrometry Goes In-Tissue. Cell Syst. 6, 10–12 (2018).

10. Ser, Z., Cifani, P. & Kentsis, A. Optimized Cross-Linking Mass Spectrometry for in Situ Interaction Proteomics. J. Proteome Res. 18, 2545–2558 (2019).

11. Jiang, P.-L. et al.. A Membrane-Permeable and Immobilized Metal Affinity Chromatography (IMAC) Enrichable Cross-Linking Reagent to Advance In Vivo Cross-Linking Mass Spectrometry. Angew. Chem. Int. Ed. 61, e202113937 (2022).

12. Iacobucci, C. et al.. First Community-Wide, Comparative Cross-Linking Mass Spectrometry Study. Anal. Chem. 91, 6953–6961 (2019).

13. Asor, R. & Kukura, P. Characterising biomolecular interactions and dynamics with mass photometry. Curr. Opin. Chem. Biol. 68, 102132 (2022).

14. Young, G. et al.. Quantitative mass imaging of single biological macromolecules. Science 360, 423–427 (2018).

15. Young, G. & Kukura, P. Interferometric Scattering Microscopy. Annu. Rev. Phys. Chem. 70, 301–322 (2019).

16. Dong, J., Maestre, D., Conrad-Billroth, C. & Juffmann, T. Fundamental bounds on the precision of iSCAT, COBRI and dark-field microscopy for 3D localization and mass photometry. J. Phys. Appl. Phys. 54, 394002 (2021).

17. Wu, D. & Piszczek, G. Standard Protocol for Mass Photometry Experiments. Eur. Biophys. J. EBJ 50, 403– 409 (2021).

18. Olerinyova, A. et al.. Mass Photometry of Membrane Proteins. Chem 7, 224–236 (2021).

19. Paul, S. S., Lyons, A., Kirchner, R. & Woodside, M. T. Quantifying Oligomer Populations in Real Time during Protein Aggregation Using Single-Molecule Mass Photometry. ACS Nano 16, 16462–16470 (2022).

20. den Boer, M. A. et al.. Comparative Analysis of Antibodies and Heavily Glycosylated Macromolecular Immune Complexes by Size-Exclusion Chromatography Multi-Angle Light Scattering, Native Charge Detection Mass Spectrometry, and Mass Photometry. Anal. Chem. 94, 892–900 (2022).

21. Sonn-Segev, A. et al.. Quantifying the heterogeneity of macromolecular machines by mass photometry. Nat. Commun. 11, 1772 (2020).

22. Foley, E. D. B., Kushwah, M. S., Young, G. & Kukura, P. Mass photometry enables label-free tracking and mass measurement of single proteins on lipid bilayers. Nat. Methods 18, 1247–1252 (2021).

23. Niebling, S. et al.. Biophysical Screening Pipeline for Cryo-EM Grid Preparation of Membrane Proteins. Front. Mol. Biosci. 9, |p(2022).

24. Lai, S.-H., Tamara, S. & Heck, A. J. R. Single-particle mass analysis of intact ribosomes by mass photometry and Orbitrap-based charge detection mass spectrometry. iScience 24, 103211 (2021).

25. Wu, D., Hwang, P., Li, T. & Piszczek, G. Rapid characterization of adeno-associated virus (AAV) gene therapy vectors by mass photometry. Gene Ther. 29, 691–697 (2022).

26. Ebberink, E. H. T. M., Ruisinger, A., Nuebel, M., Thomann, M. & Heck, A. J. R. Assessing production variability in empty and filled adeno-associated viruses by single molecule mass analyses. Mol. Ther. - Methods Clin. Dev. 27, 491–501 (2022).

27. Lim, W. K., Rösgen, J. & Englander, S. W. Urea, but not guanidinium, destabilizes proteins by forming hydrogen bonds to the peptide group. Proc. Natl. Acad. Sci. 106, 2595–2600 (2009).

28. Das, A. & Mukhopadhyay, C. Urea-Mediated Protein Denaturation: A Consensus View. J. Phys. Chem. B 113, 12816–12824 (2009).

29. Huerta-Viga, A. & Woutersen, S. Protein Denaturation with Guanidinium: A 2D-IR Study. J. Phys. Chem. Lett. 4, 3397–3401 (2013).

30. Steigenberger, B., Pieters, R. J., Heck, A. J. R. & Scheltema, R. A. PhoX: An IMAC-Enrichable Cross-Linking Reagent. ACS Cent. Sci. 5, 1514–1522 (2019).

31. Gorynia, S. et al.. Structural and functional insights into a dodecameric molecular machine – The RuvBL1/RuvBL2 complex. J. Struct. Biol. 176, 279–291 (2011).

32. Maurizy, C. et al.. The RPAP3-Cterminal domain identifies R2TP-like quaternary chaperones. Nat. Commun. 9, 2093 (2018).

33. Seraphim, T. V. et al.. Assembly principles of the human R2TP chaperone complex reveal the presence of R2T and R2P complexes. Structure 30, 156-171.e12 (2022).

34. Chen, F., Nielsen, S. & Zenobi, R. Understanding chemical reactivity for homo- and heterobifunctional protein cross-linking agents: Chemical cross-linking efficiency in proteins. J. Mass Spectrom. 48, 807–812 (2013).

35. Beveridge, R., Stadlmann, J., Penninger, J. M. & Mechtler, K. A synthetic peptide library for benchmarking crosslinking-mass spectrometry search engines for proteins and protein complexes. Nat. Commun. 11, 742 (2020).

36. Ihling, C. H., Piersimoni, L., Kipping, M. & Sinz, A. Cross-Linking/Mass Spectrometry Combined with Ion Mobility on a timsTOF Pro Instrument for Structural Proteomics. Anal. Chem. 93, 11442–11450 (2021).

37. Perez-Riverol, Y. et al.. The PRIDE database resources in 2022: a hub for mass spectrometry-based proteomics evidences. Nucleic Acids Res. 50, D543–D552 (2022).

