## Supplementary material for "Denaturing mass photometry for straightforward optimization of protein-protein cross-linking reactions at single-molecule level": supp data all

**Reagents.**

2-β mercaptoethanol (M3148), Ammonium sulfate (A-4418), Brilliant Blue G250 (B-0770), Hepes (H3375), Sodium Chloride (NaCl, 59625), Phosporic acid (7664-38-2), Sodium dodecyl sulfate (SDS, S7920), Trizma base (Tris HCl, T-5941) were purchased from Sigma-Aldrich. Glycerol (BP229-1) and Methanol (M/4062/17) were purchased from Fischer Scientific. Dimethylsulfoxide was purchased from Merck. Absolute anhydrous ethanol was purchased from Carlo Erba. Nitrogen was purchased from Linde. Milli-Q water was generated using an Arium® Pure DI Ultrapure Water System (Sartorius).

**R2SP expression and purification.**

RuvBL1ΔT127-E233 carries an N-terminal 6x His-tag followed by thrombin cleavage site, while RuvBL2ΔE134-E237 carries an C-terminal Flag+FH8 tag preceded by HRV-3C cleavage site (Flag was used for detection). The RuvBL1ΔT127-E233- RuvBL2 ΔE134-E237 complex was expressed in E. coli (DE3) (Novagen, 71400), with 100 µM IPTG overnight at 18 °C. The complex was immobilized in a 5 ml HistrapTM HP (GE Healthcare) and eluted with 300mM imidazole. Peak fractions collected from the HisTrap were incubated with 5mM CaCl2 during 1 h and loaded onto an HiPrepTM Octyl FF 16/10 column (GE Healthcare). Bound proteins were eluted using 5mM EDTA. To remove the FLAG_FH8 tag the collected samples were incubated 18 h at 4 °C with 1 % (w/w) HRV-3C protease (Thermo Fisher Scientific). A final Superose 6 equilibrated in buffer 20 mM Tris–HCl pH 8.0, 150mM NaCl, 5% glycerol, 2mM MgCl2 and 0.5mM TCEP, was used to separate a stable dodecameric peak, from the HRV3C protease and cleaved tags. The pooled dodecamer was concentrated to 37.75 mg/ml using a 30 kDa Cut-off Amicon Ultra centrifugal filter (Millipore).

SPAG1(622-926) carries an C-terminal Flag-tag without a cleavage site, while PIH1D2(231-315) has a C-terminal StrepTag II preceded by a Human Rhino Virus 3C cleavage site (HRV-3C). SPAG1(622-926)/PIH1D2(231-315) were co-expressed in Escherichia coli (DE3*) (Novagen, 71400), with 50 µM IPTG overnight at 18 °C in a New Brunswick™ (Innova®) 44R Shaker at 150 rpm. The SP_mini complex was immobilized in a 5 ml StrepTactin XT HC (IBA life sciences), and eluted with 50 mM Biotin. Peak fractions collected from the StrepTactin XT were Injected in a Superdex 200 16/60 XK equilibrated in Buffer 20 mM Hepes pH 8, 300 mM NaCl, 0.5 mM TCEP, allowing the isolation of a heterodimer. Collected fractions from the main peak were diluted to reduce the concentration of NaCl to 50 mM, and further polished in ResourceTM Q (GE), and eluted with a linear gradient, allowing the separation of a major peak corresponding to the intact complex (w/o degradation) at approximately 170 mM NaCl. Collected peak fractions were supplemented with 20 mM imidazole and tag removal was performed by incubating 18 h at 4 °C with 1% (w/w) HRV-3C protease (Thermo Fisher Scientific). Digested sample was injected in a 5ml StrepTactin XT in tandem with 1ml HisTrap. The collected flow was concentrated to 14.7 mg/ml using a 3 kDa Cut-off Amicon Ultra centrifugal filter (Millipore). All purification steps were carried out at room temperature and were monitored by NuPAGE Bis-Tris gels (Invitrogen, NP0302).

**R2SP complex formation.**

R2SP complex was formed by mixing pure RuvBL1(ΔT127-E233)/RuvBL2(ΔE134-E237) with excess pure SPAG1(622-926)/PIH1D2(231-315) complex at a ratio of 1 : 4 (considering RuvBLs dodecameric, and SPAG1/PIH1D2 heterodimeric) over night at 4 °C. Formed R2SP complex was separated from free excess SPAG1/PIH1D2 using a Superose 6 16/60 XK (GE Healthcare) previously equilibrated in 20 mM Hepes pH 8, 150 mM NaCl, 2mM MgCl2, and 0.5 mM TCEP. The eluted peak was concentrated to 7.5 mg/ml using a 3 kDa Cut-off Amicon Ultra centrifugal filter (Millipore)

**Cross-linking mass spectrometry.**

R2SP complex was cross-linked in triplicates with 25, 100, 400 molar excesses of DSBU (45 min, 18°C), as described in Methods part, before quenching reaction with 15 mM Tris HCl (20 min). Samples were reduced by adding DTT to a final concentration of 5 mM and incubation at 37°C for 30 min. The alkylation was done by adding Iodoacetamide to a final concentration of 15 mM (1 hour incubation step in the dark). Samples were then processed with overnight digestion with Trypsin/Lys-C (Promega, Madison, USA) at a 50:1 substrate:enzyme ratio (w/w) at 37°C overnight. The digestions were finally quenched with 1 % TFA.

Peptides were cleaned up by using the AssayMAP Bravo platform (Agilent Technologies; Santa Clara, California) with 5µL C18 cartridges (Agilent). Cartridges were primed with 100 µl 0.1 % TFA in 80 % ACN and equilibrated with 50 µl 0.1 % TFA in H2O. 180 µl of digested peptides diluted in equilibration buffer were loaded on the cartridges and washed with 50 µl equilibration buffer. Peptides were eluted with 50 µl 0.1 % TFA in 80 % CAN and stored at -80°C prior to the mass spectrometry analysis. Cross-linked peptides were dried in a SpeedVac concentrator and resuspended in 10 µl of 2 % ACN/0.1 % formic acid.

NanoLC-MS/MS analysis was performed using a nanoAcquity UPLC (Waters, Milford, USA) hyphenated to a Q Exactive HF-X mass spectrometer (Thermo Fisher Scientific, Bremen, Germany) equipped with a nanoSpray source. After trapping on a NanoEase M/Z Symmetry pre- column (C18, 100 Å, 5 µm, 180 µm × 20 mm; Waters), samples were separated on a NanoEase M/Z BEH column (C18, 130 Å, 1.7 µm, 75 µm x 250 mm; Waters) maintained at 60°C. A gradient of 102 min was applied: mobile phases A (0.1% v/v formic acid in H2O) and B (0.1% v/v formic acid in ACN). The following conditions were applied: 3 % B for 3 min, 3–40 % B for 90 min, – 90% B for 1 min, 90% B for 5 min, 90–1% B for 2 min and finally 1% B maintained for 2 min (flow rate of 350 nl/min). Acquisition in Data Dependant Acquisition mode (Top 10 precursor ions) was done using following parameters: MS resolution of 120.000 (AGC target 3e6), MS/MS resolution of 30.000 (AGC target of 2e5), 3-7 charge states enable, HCD stepped collision energy (27, 30, 33 % normalized collision energy). Raw data were directly processed with Thermo Proteome Discoverer 2.5.0.400 (Thermo Scientific) using the XlinkX node for identification of crosslinks and the Sequest HT node for the identification of linear peptides. For both linear and cross-linked peptides searches, Cystein carbamidomethylation was set as fixed modification. Methionine oxidation, N-term acetylation, tris-quenched mono-links and water-quenched mono-links were set as dynamic modifications. Trypsin was set as the cleavage enzymes with minimal length of 7 amino acids, 2 (linear peptides) and 3 (cross-linked peptides) missed cleavages were allowed. To increase confidence, identification were only accepted for Maximal XlinkX scores > 40. A 1% false discovery rate was applied at the cross-linked peptides level (XlinkX validator node). For the consensus step, proteins identifications were controlled at a 1% FDR in the protein validator node. Out of the three replicates performed for each molar excess of DSBU, unique cross-link sites were validated when present in at least 2 out of 3 replicates.

**Supplementary Figures.**

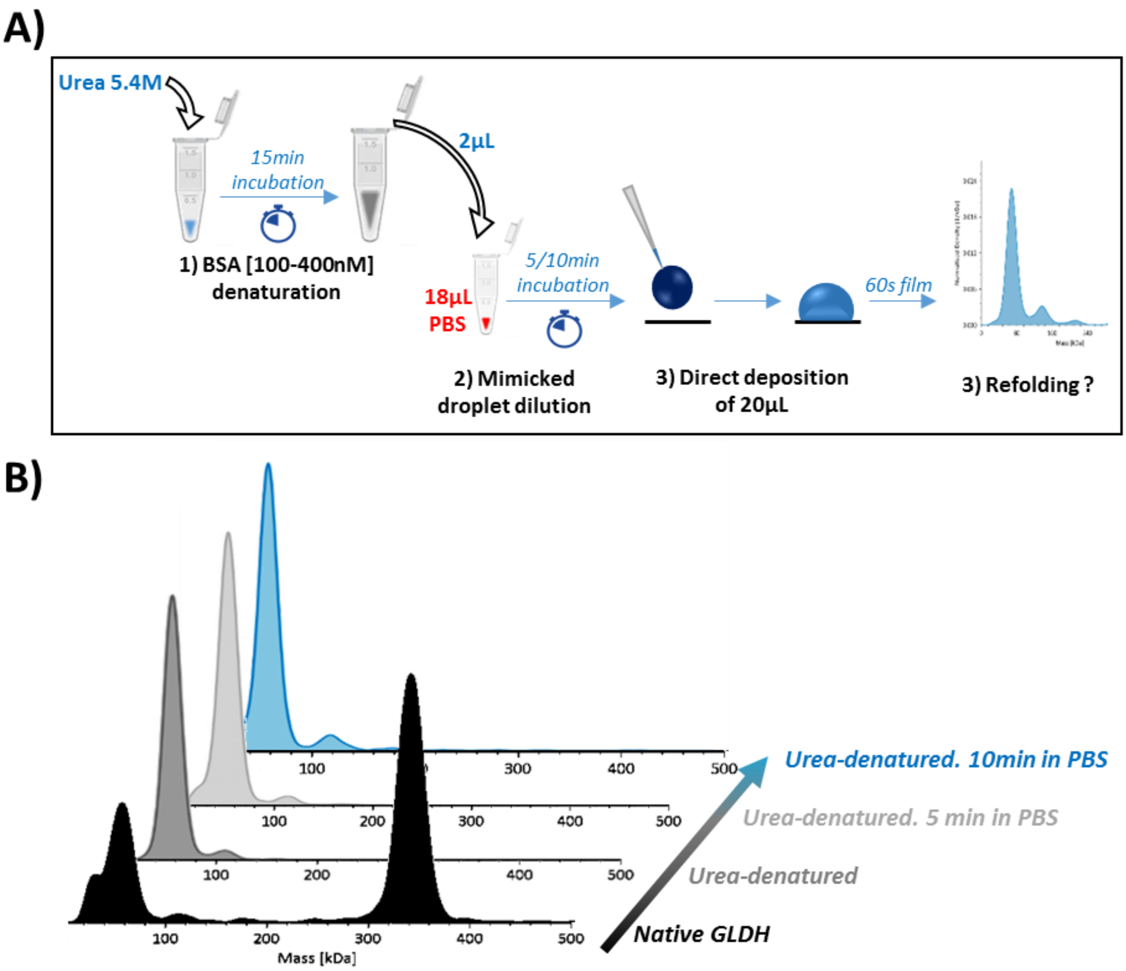


**Figure S1. GLDH refolding study results mimicking a PBS droplet dilution. A)** GLDH has been first denatured, then diluted to the tenth in a PBS tube. After 0 min, 5 min and 10 min this solution was directly analyzed in MP using the buffer-free focusing mode. **B)** MP profiles of native GLDH, GLDH right after denaturation and after 5 and 10 min dilution in PBS.

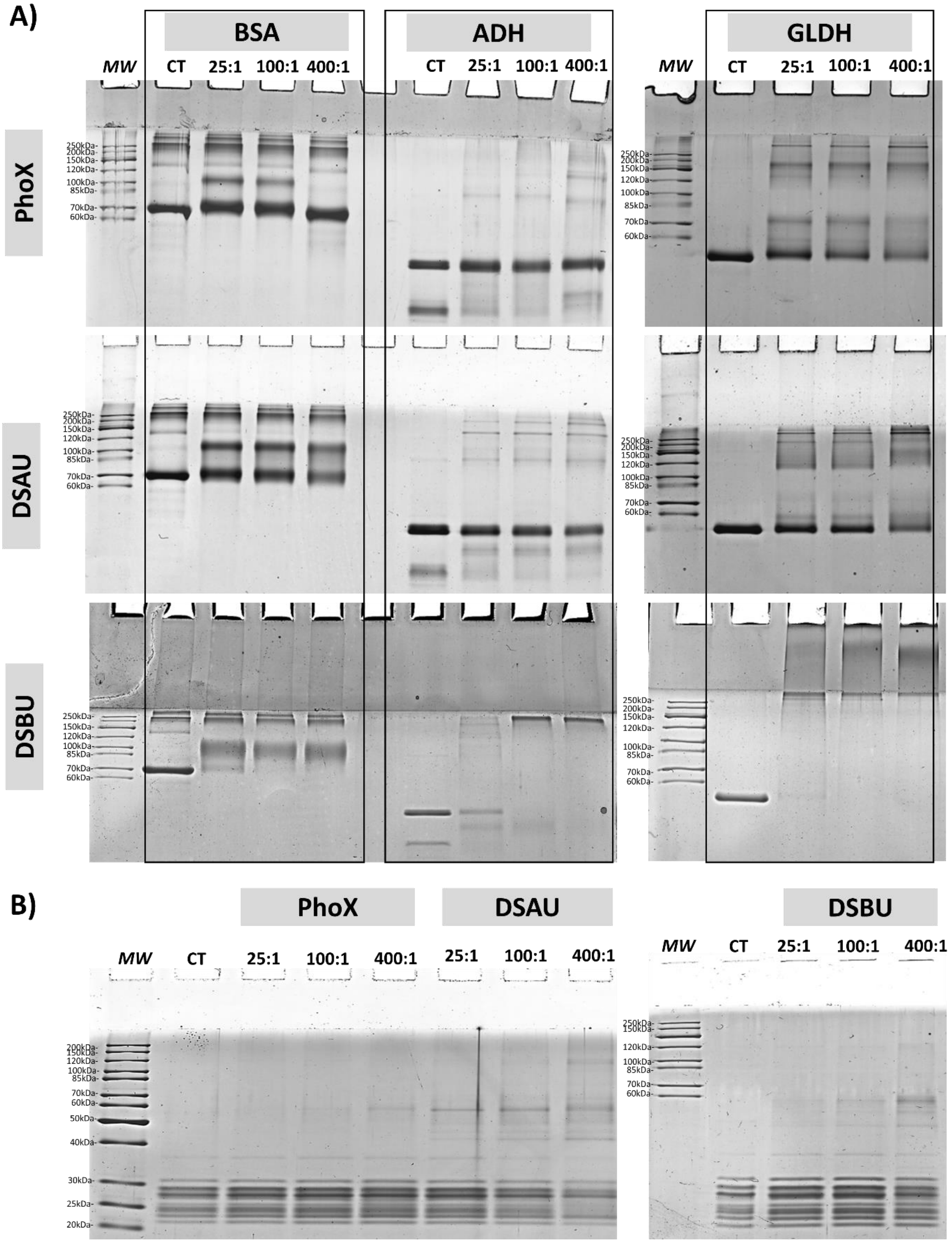


**Figure S2. SDS-PAGE migrations of A)** BSA, ADH, GLDH cross-linked with 25, 100, 400 molar excesses of PhoX, DSAU, DSBU**. B)** 20S proteasome cross-linked with 25, 100, 400 molar excesses of PhoX, DSAU, DSBU.

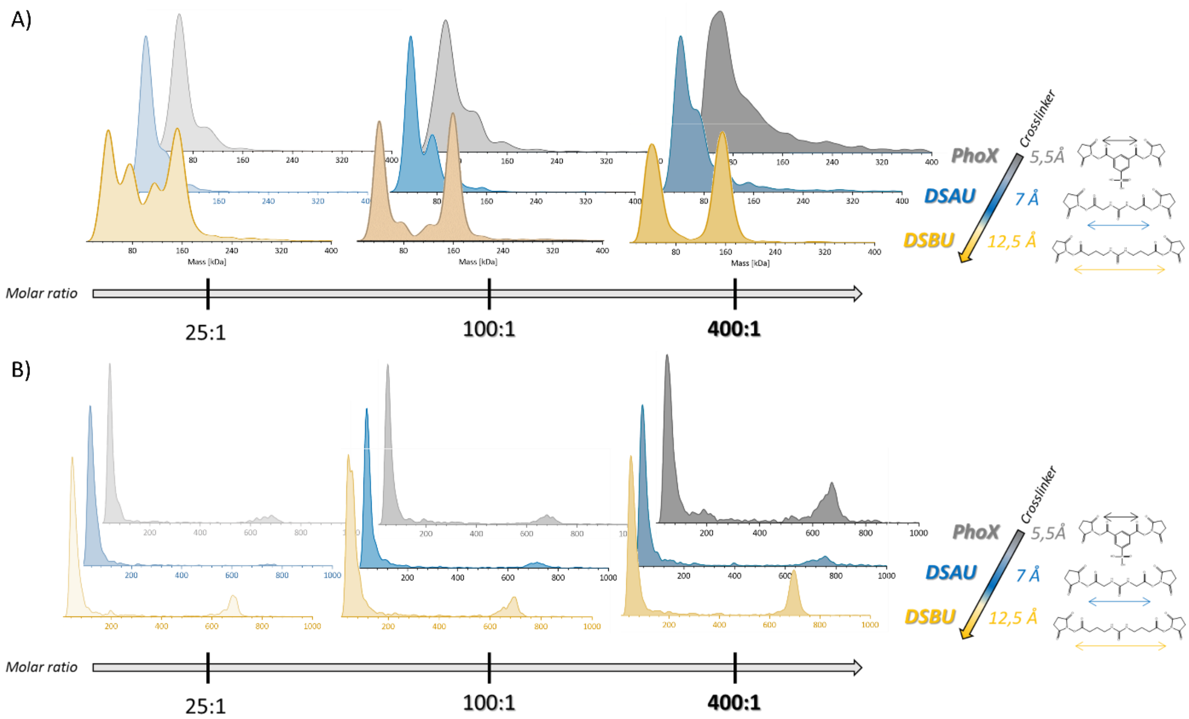


**Figure S3. Effect of cross-linking reagent (size, flexibility) on oligomeric states stabilized, measured in dMP: presented results are probability densities (KD) of A)** ADH and **B)** 20S proteasome, cross-linked with increasing molar ratios of PhoX, DSAU, DSBU.

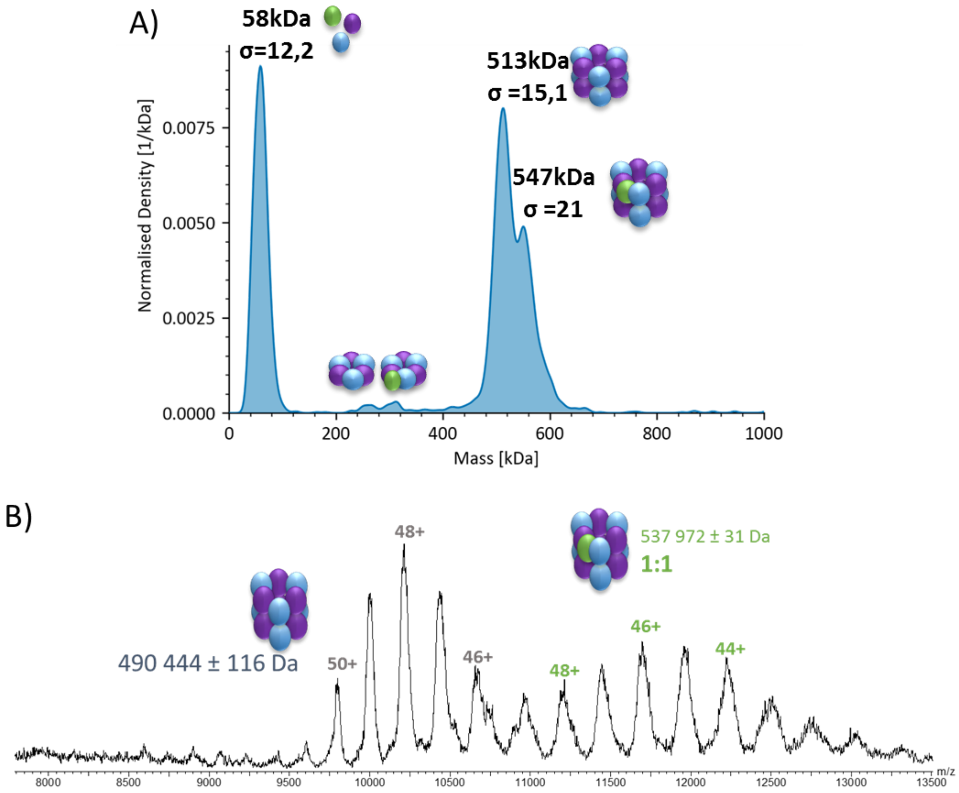


**Figure S4. Oligomeric states of R2SP complex A)** nMP profile of R2SP complex represented as Kernel Density**.** **B)** Native-MS spectrum of R2SP complex.

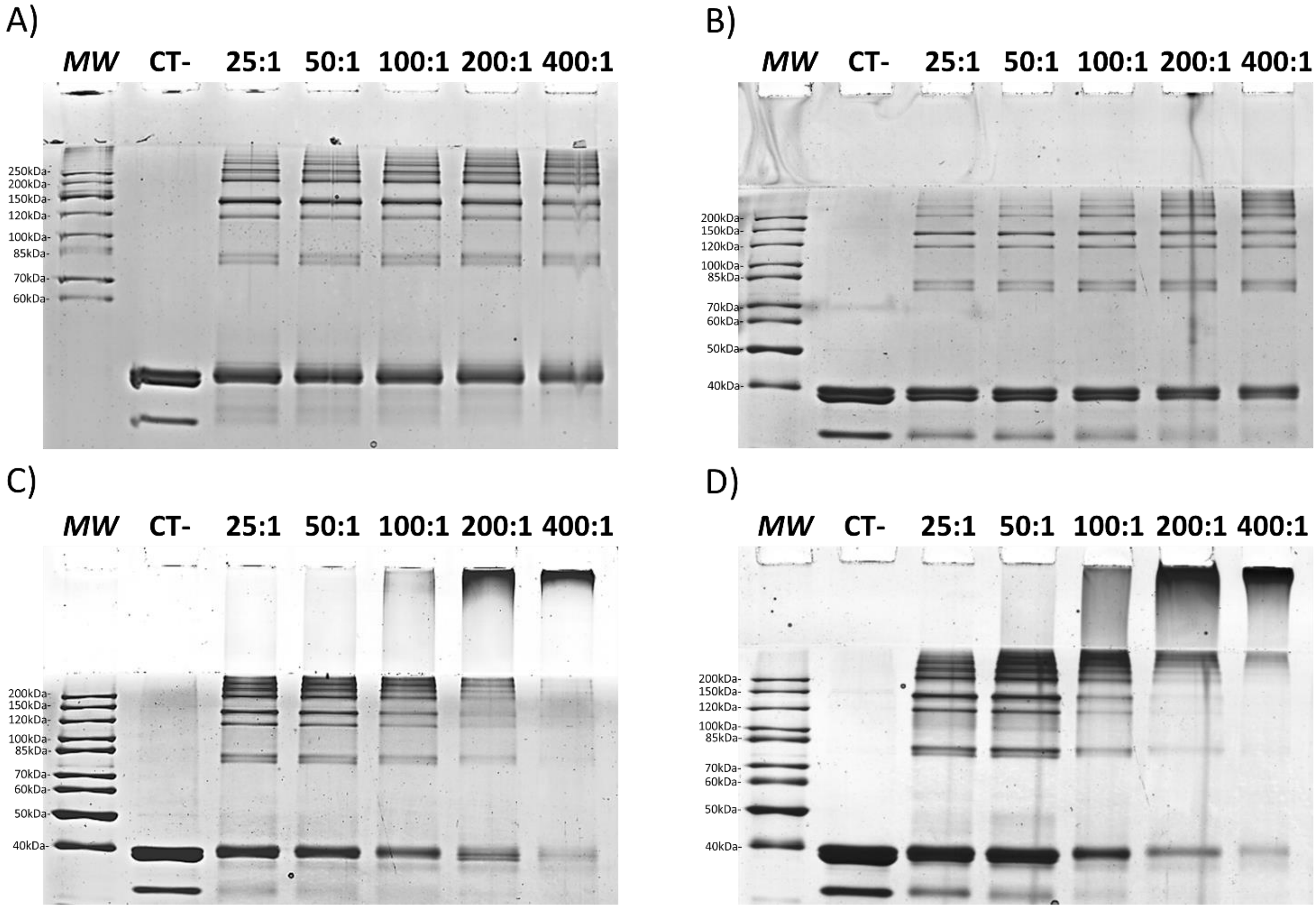


**Figure S5. SDS-PAGE of R2SP complex cross-linked with 25, 50, 100, 200, 400 molar excesses of A)** PhoX**, B)** DSAU**, C)** DSSO**, D)** DSBU**.**

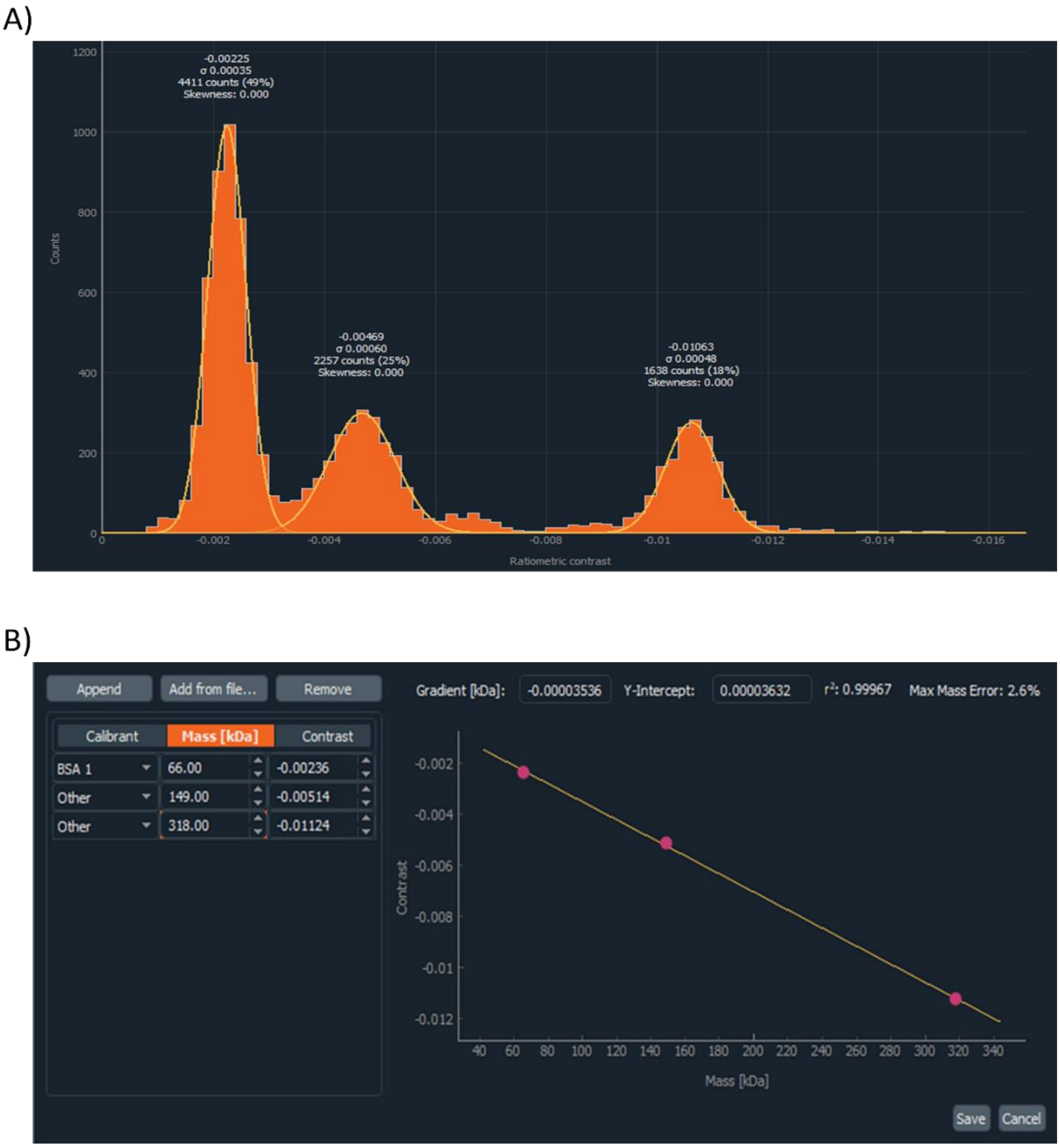


**Figure S6. A) Example of MP mass histogram integration of a mix BSA/Bevacizumab/GLDH.** Peak have been integrated at their half-height. **B)** Example of contrast-to-mass calibration using the mix BSA/Bevacizumab/GLDH.

**Supplementary Tables.**

**Table S1. Protein-free droplet measurements with decreasing denaturant concentrations showing the impact of denaturant droplet concentration on Sharpness, Brightness and Signal parameters.**

| | Droplet concentration (M) | Sharpness (%) | Brightness (%) | Signal (%) |
|---|---|---|---|---|
| PBS | **2.9** | **5** | **73** | **0.03** |
| Urea | **5.4** | **3** | **44** | **0.05** |
| | **2.5** | **3** | **55** | **0.03** |
| | **1.75** | **5** | **63** | **0.03** |
| | **0.8** | **7** | **67** | **0.03** |
| Guanidine | **5.4** | **1** | **29** | **0.05** |
| | **2.5** | **3** | **47** | **0.04** |
| | **1.75** | **4** | **60** | **0.03** |
| | **0.8** | **4** | **62** | **0.03** |
| | **0.4** | **5** | **67** | **0.03** |

**Table S2. Mass measurements and counts number after nMP and dMP measurements of BSA.**

| | | PBS | Urea 5,4M | | Guanidine HCl 6M | |
|---|---|---|---|---|---|---|
| | | | 2h | 16h | 2h | 16h |
| μ of gaussian fit (kDa) | Monomer (66,4 kDa) | **75,0 ± 2,6** | **73,3 ± 1,5** | **71,7 ± 1,2** | **72,7 ± 2,1** | **73,7 ± 0,6** |
| | Dimer (133 kDa) | **144,3 ± 3,8** | **147,0 ± 4,4** | **142,3 ± 1,2** | **145,0 ± 3,5** | **142,3 ± 5,7** |
| σ of gaussian fit (kDa) | Monomer (66,5 kDa) | **8,2 ± 1,8** | **9,1 ± 0,9** | **7,4 ± 1,1** | **10,2 ± 1,4** | **11,7 ± 1,6** |
| | Dimer (133 kDa) | **8,4 ± 1,3** | **14,5 ± 1,4** | **8,1 ± 2,3** | **13,3 ± 0,8** | **13,5 ± 2,4** |

**Table S3. Statistical results after XL-MS of R2SP cross-linked in triplicates at 25, 100, 400 molar excesses of DSBU.** Only validated unique cross-links (present in at least 2 out of 3 replicates) have been taken into account. Unique XL have been differentiated between intra- and inter-protein XL.

| | **25:1** | **100:1** | **400:1** |
|---|---|---|---|
| Validated unique XL (2/3) | 94 | 97 | 71 |
| % intra-XL | 55 % | 53 % | 46 % |
| % inter-XL | **45 %** | **47 %** | **54%** |

**Table S4. Example of $Eff_{XL}$ calculation on R2SP complex cross-linked with 400 molar excesses of DSBU**.

| **Measurement replicate** | **Monomer Counts** | **Total counts** | **% of oligomeric states >1** | **$Eff_{XL}$** |
|---|---|---|---|---|
| 1 | 528 | 2006 | $\frac{2006-528}{2006} * 100 = 74\ \%$ | **Mean = 77 % ± 5%** |
| 2 | 892 | 5026 | $\frac{5026-892}{5026} * 100 = 82\ \%$ | |
| 3 | 690 | 2735 | $\frac{2735-690}{2735} * 100 = 75\ \%$ | |

**Table S5. Example of $SF_{XL}$ calculation on R2SP complex cross-linked with 400 molar excesses of DSBU.**

| | R2SP abundance (%) | $SF_{XL}$ |
|---|---|---|
| **nMP** of R2SP | 46 | $\frac{0,27}{0,46} = \mathbf{0,6}$ |
| **dMP** of 400:1 DSBU:R2SP | 27 | |
